# End-to-End Differentiable Blind Tip Reconstruction for Noisy Atomic Force Microscopy Images

**DOI:** 10.1101/2022.09.24.509314

**Authors:** Yasuhiro Matsunaga, Sotaro Fuchigami, Tomonori Ogane, Shoji Takada

## Abstract

Observing the structural dynamics of biomolecules is vital to deepening our understanding of biomolecular functions. High-speed (HS) atomic force microscopy (AFM) is a powerful method to measure biomolecular behavior at near physiological conditions. In the AFM, measured image profiles on a molecular surface are distorted by the tip shape through the interactions between the tip and molecule. Once the tip shape is known, AFM images can be approximately deconvolved to reconstruct the surface geometry of the sample molecule. Thus, knowing the correct tip shape is an important issue in the AFM image analysis. The blind tip reconstruction (BTR) method developed by Villarrubia^1^ is an algorithm that estimates tip shape only from AFM images using mathematical morphology operators. While the BTR works perfectly for noise-free AFM images, the algorithm is susceptible to noise, or it is difficult to determine a threshold parameter against noise. To overcome this issue, we here propose an alternative BTR method, called *end-to-end differentiable* BTR, based on a modern machine learning approach. In the method, we introduce a loss function and a regularization term to prevent overfitting to noise, and the tip shape is optimized with automatic differentiation and backpropagations developed in deep learning frameworks. Using noisy pseudo-AFM images of myosin V motor domain as test cases, we show that our end-to-end differentiable BTR is robust against noise in AFM images. The method can also detect a double-tip shape and deconvolve doubled molecular images. Finally, application to real HS-AFM data of myosin V walking on an actin filament shows that the method can reconstruct the accurate surface geometry of actomyosin consistent with the structural model. Our method serves as a general post-processing for reconstructing hidden molecular surfaces from any AFM images. Codes are available at https://github.com/matsunagalab/differentiable_BTR.

## Introduction

Atomic force microscopy (AFM) is a unique technique for imaging structures of sample molecules bound to a surface at ambient conditions ^2^. Recently, high-speed AFM (HS-AFM) with dramatically faster imaging rates (up to tens of frames per second) has been developed, enabling us to directly observe biomolecules in action^3 4^. HS-AFM can investigate detailed structure-function relationships in biomolecules that cannot be observed with other methods, and it has been established as one of indispensable techniques in modern biophysics. Example applications of HS-AFM include myosin V walking along an actin filament^5^, rotary catalysis of F_1_-ATPase^6^, structural dynamic of intrinsically disordered protein^7^, and the functional dynamics of CRISPR-Cas9 in action^8^. Currently, the spatial resolutions of HS-AFM instruments are ∼ 2 nm in the lateral direction and ∼0.15 nm in the vertical direction to the AFM stage^9^.

Importantly, separately from the resolution of the image profile, the resolutions of the obtained sample surface information are further limited by the tip geometry and the tip-sample interactions. The relationship among the tip geometry, the image profile, and the sample surface is shown in Figs. 1a and 1b. When the tip is sufficiently thin, the obtained image profile is nearly the same as the sample surface (Fig. 1a). On the other hand, when the tip is blunt compared to the scale of samples, the image profile is blurred from the sample surface (Fig. 1b). Once the tip shape is known, algorithms, called *erosion*, have been proposed to “deconvolve” the image profile for reconstructing approximate sample surface geometry.^1 10 11^ Thus, to reconstruct the surface geometry of the sample molecule, it is crucial to know the tip shape accurately. Tip shape estimation is also important for inferring 3D molecular structures from AFM images. In the recent analysis of AFM images, pseudo-AFM images are emulated from 3D molecular structures (obtained with different experimental or computational techniques, e.g., X-ray crystallography and molecular dynamics simulations) and an assumed tip shape and then compared with the experimental AFM image ^12 13 14 15 16 17 18 19 20 21^. In the analysis, the 3D structure which generates the pseudo-AFM images most similar to the experimental AFM image is selected as the best estimate for 3D molecular structure. In this kind of analysis, the accuracy of pseudo-AFM images crucially depends on the tip shape. Thus, the tip shape should be determined as accurately as possible. Currently, there are three possible approaches to obtain a tip shape^22 23^. The first approach is the direct imaging of a tip using either a scanning or transmission electron microscope (SEM or TEM). However, both SEM and TEM provide only a two-dimensional projection of the sample. It is unrealistic to install special equipment for imaging the tip from various angles as a routine for determining the tip shape. Moreover, as the tip can be damaged over time during AFM measurement, determining tip shape before or after the AFM measurement is not necessarily appropriate. The second approach is to estimate the tip shape during the AFM measurement. This approach determines the tip shape from samples whose geometry is *a priori* known^24 25 26^. With the combination of mathematical modeling, an approximate tip shape can be reconstructed. For example, Niina et al. recently proposed a method to estimate a tip shape by comparing the pseudo-AFM images generated from 3D molecular structures with AFM images^20^. By assuming that the tip shape is a hemisphere (radius *r*) combined with a circular frustum of a cone (half angle *θ*), they proposed to estimate these two geometric parameters. However, since real tips can be of any shape, e.g., double tips with two separate acutes, the assumption of a shape limits the application of the method.

**Figure 1.**
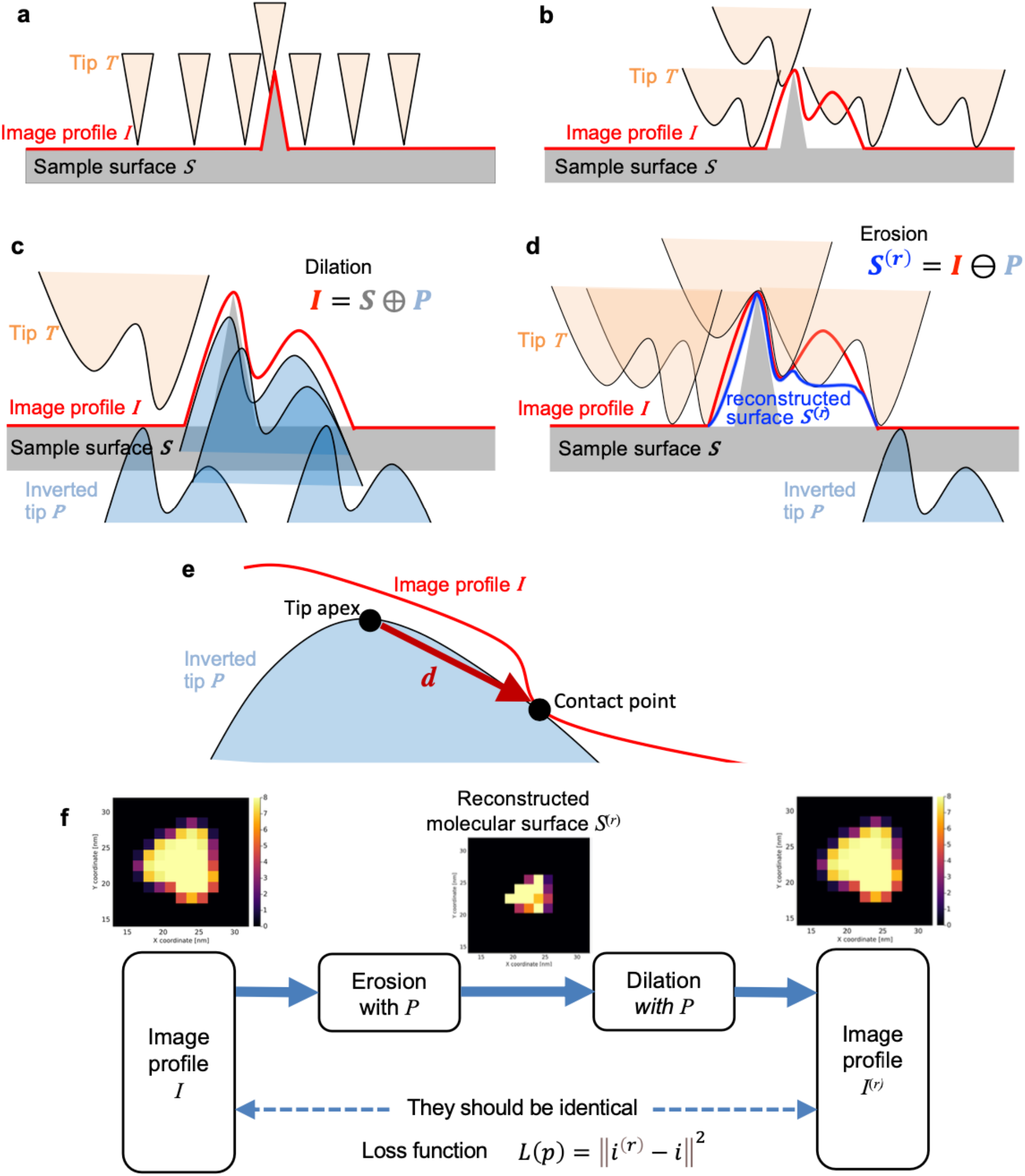
Morphology operators and schematic of differentiable blind tip reconstruction (BTR). **a**, Image profile obtained by scanning a surface with a thin tip. **b**, Image profile obtained by scanning the same surface as (**a**) with a blunt tip. **c**, Schematic picture illustrating a morphology operator, *dilation*. **d**, Schematic picture illustrating a morphology operator, *erosion*. Here, for the sake of intuitive explanation, erosion is represented with a non-inverted tip instead of its inversion. **e**, Relation of a contact point and tip apex position, explaining the idea behind the original BTR. **f**, Schematic of differentiable BTR.

The third approach estimates the tip shape through the image analysis of AFM data without any prior knowledge of sample molecule and tip shape. The blind tip reconstruction^1^ developed by Villarrubia is a classical method to estimate arbitrary tip shapes from AFM images. The idea behind the algorithm is to recognize that sizes and depths of dents in the image profile must be smoother than the sharpness of the tip. Then, the algorithm “carves” an initially blunt tip according to the sizes and depths of dents (details will be described below). While the BTR works perfectly for noise-free AFM images, its algorithm is susceptible to noise, or it is difficult to determine a threshold parameter against noise. This is a crucial issue for the analysis of HS-AFM images because HS-AFM is generally more prone to noise than conventional AFM. Over these two decades, several improvements or new methods have been proposed for better estimation of tip shape; Dongmo et al. proposed monitoring tip volume for tuning *thresh* parameter for noisy AFM images. Tian et al. extended Villarrubia’s original BTR with the dexel representation for reconstructing general 3D tip shapes and proposed an improved regularization scheme against noise^27^. Jóźwiak et al. simplified Tian’s idea to the case of standard AFM tips^28^. Flater et al. proposed a systematic way to determine the threshold parameter against noisy AFM images^22^. By approximating mathematical morphology operators by linear operators, Bakucz et al. proposed a reconstruction method based on the Expectation-Maximization (EM) algorithm with the tip shape represented as a hidden variable^29^. Despite these studies, the original BTR is not routinely utilized in the analysis of noisy AFM data, especially for the analysis of HS-AFM data, due to its susceptibility to noise and difficulty in tuning the parameter.

The reason why Villarrubia’s BTR is susceptible to noise is that it is difficult to correctly determine whether the tip shape geometry or noise causes an individual dent in the image profile. In the algorithm of the BTR, once a dent caused by noise is misinterpreted as the cause of the geometry of the tip, the tip is “carved” to be fitted to the noise. This leads to a prediction of very thin tip shape. In terms of machine learning theory, this can be regarded as overfitting the tip shape to noise.

To prevent such overfitting, we here introduce an appropriate loss function and a regularization term considering noise statistics, which are absent in the original BTR. As the morphology operators (Figs 1c and 1d) used in the loss function are complicated nonlinear functions, minimizing such a complicated loss function over tip shape is challenging. The technologies developed by recent advances in deep learning studies^30^ can potentially overcome this problem. Automatic differentiation and smart optimizers are recently being applied to optimize not only the parameters of neural networks but also the parameters of physical models, by implementing physical functions as differentiable functions^31 32^. For example, Zhou et al. recently modeled the Lorentz TEM observation process as a differentiable neural network layer and successfully solved the inverse problem of phase retrieval stably by backpropagation^33^. In the current study, we propose to use these technologies to optimize the loss function over the tip shape. Our method (called the end-to-end differentiable BTR) implements morphology operators as differentiable functions and optimizes the loss function in the same way as neural networks under a neural network framework. Using pseudo-AFM images generated from a known molecular structure (myosin V motor domain) as a test case, we show that the differentiable BTR is robust against noise in AFM images, as well as lower parameter dependence compared to the original BTR. Furthermore, the differentiable BTR, but not the original BTR, can correctly detect double tips, one of the artifacts that frequently occur in AFM measurements. Finally, we applied the method to real noisy HS-AFM images, myosin V walking along an actin filament.

## Results

We briefly review the original BTR by Villarrubia^1^, then introduce the end-to-end differentiable BTR. First, let the AFM stage be the *xy*-plane, and the height perpendicular to the stage be *z*-coordinate, where the stage position corresponds to *z* = 0. Then, the height of the sample surface at coordinates (*x, y*) is denoted by *s*(*x, y*) (Figs. 1a and 1b). The image profile obtained by scanning the sample surface with a tip is denoted by *i*(*x, y*). Then, the coordinate system for tip shape is introduced. We define the *uv*-plane parallel to the *xy*-plane, and its origin is located at the top of the tip apex, i.e., the tip apex corresponds to *z* = 0 in the *uv*-coordinate system. The *uv*-coordinate system is a moving coordinate that moves with the tip. The height of the tip surface is represented by *t*(*u, v*) ≥ 0, which determines the tip shape. For convenience, let us consider an inversion of the tip shape through the origin and define the surface of the inverted tip as *p*(*u, v*) = −*t*(−*u*, −*v*) ≤ 0. Then, the image profile *i*(*x, y*) can be obtained using *dilation*, which is an operator in the mathematical morphology^1^,

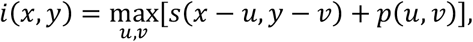

Conceptually, this would be viewed as a “convolution” of the molecular surface *s*(*x, y*) with a kernel function *p*(*u, v*) (the inverted tip shape) (Fig. 1c). But note that the dilation is not the same as convolution because convolution is a linear transformation while dilation is a nonlinear transformation, representing the physical interaction between tip and sample. Following the terminology of mathematical morphology, let us denote the sets of all points on or below the surfaces as *umbras*, and three umbras for surface *s*(*x, y*), *i*(*x, y*), and *i*(*x, y*) defined by *S* = {(*x, y, z*) | *z* ≤ *s*(*x, y*)}, *I* = {(*x, y, z*) | *z* ≤ *i*(*x, y*)}, and *P* = {(*u, v, z*) | *z* ≤ *p*(*u, v*)}, respectively. Then the dilation can be simply written as *I* = *S* ⊕ *P*. Given the umbras *I* of a measured image profile *i*(*x, y*) and an inverted tip shape *P*, the sample surface *S* can be approximately reconstructed by *erosion*, written as *S*^(*r*)^ = *I* ⊖ *P*, which is another operator in the mathematical morphology defined by,

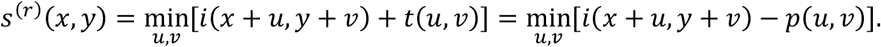

Conceptually, this would be viewed as a “deconvolution” that “carves” the image profile with a kernel function *p*(*u, v*) (Fig. 1d). But note again that the erosion is a nonlinear transformation. From the properties of the erosion, we can say that *S*^(*r*)^ ⊕ *P* = (*I* ⊖ *P*) ⊕ *P* = *I* and *S*^(*r*)^ ⊇ *S*, which means that *S*^(*r*)^ is the (least) upper bound on sample surfaces that reproduces the image profile *I* with the tip shape *P*. The dilation of the erosion of A by B is called *opening*, and is written as (*A* ⊖ *B*) ⊕ *B* = *A* ∘ *B*, so the above relation can be written simply as *I* ∘ *P* = *I*.

The algorithm of Villarrubia’s BTR (hereafter called the *original* BTR) estimates an upper bound on the tip shape from AFM images. The original BTR starts from the relation, *I* ∘ *P* = *I*, and constructs an upper bound condition on *P* from this relation. An important property of opening is *I* ∘ *P* = ⋃{*P* + ***a*** | *P* + ***a*** ⊂ *I*} where ***a*** is a three-dimensional translation^34^. Combined with the relation *I* ∘ *P* = *I*, we can say *I* = ⋃{*P* + ***a*** | *P* + ***a*** ⊂ *I*}. This means that every point of *I* is contained in one or more translates of *P* while the translates are limited by the requirement that no part of the translated inverted tip extends above the surface of *I*. In particular, every point on the surface of *I* must be touched by the surface of one or more of these translates of *P*. Suppose that the translated inverted tip touches the surface of *I* at ***c*** ∈ *I*, and the vector from the origin (tip apex) of the moving *uv*-coordinates to ***c*** is defined as ***d*** (Fig. 1e). Then, the translated inverted tip can be written as *P* + ***c*** − ***d***, and the upper bound condition on the inverted tip shape is that the translated inverted tip is entirely on or below the surface of *I*, that is,

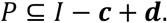

This condition holds for any ***c*** on the surface or inside of *I* where ***d*** exists.

The original BTR finds *P* that satisfies this upper bound condition. The algorithm starts from a thick tip shape (typically, a square pillar with a flat top), as an initial condition, that includes the true *P* as a subset. When the tip touches on the surface of image profile umbra *I* at point *i*(*c*_*x*_, *c*_*y*_), the height of the image profile in terms of the moving *uv*-coordinate is *i*(*c*_*x*_, *c*_*y*_) = *p*(*d*_*u*_, *d*_*v*_) at ***d***, and the heights of the image profile at other (*u, v*) locations are *i*(*c*_*x*_ + *u* − *d*_*u*_, *c*_*y*_ + *y* − *d*_*v*_). By adding the height of the tip at the touch point (*d*_*u*_, *d*_*y*_) to the difference between the two heights above, we get *dill* = *i*(*c*_*x*_ + *u* − *d*_*u*_, *c*_*y*_ + *y* − *d*_*v*_) − *i*(*c*_*x*_, *c*_*y*_) + *p*(*d*_*u*_, *d*_*v*_). By calculating *dill* for all possible ***d***, we find the minimum value of *dill*, written as *dill*_*min*_. If *dill*_*min*_ + *thresh* < *p*(*u, v*), then the tip shape is updated (or “carved”) with a smaller height, *p*(*u, v*) = *dill*_*min*_ + *thresh*. Here, *thresh* > 0 is a parameter that determines a tolerance of inconsistency between the image profile and the tip estimate. The above calculations are performed for all (*c*_*x*_, *c*_*y*_) positions on the image profile *i*(*c*_*x*_, *c*_*y*_). All the computational steps are repeated until the tip shape is converged. In the end, the upper bound on the true shape *P, P*^(*r*)^ ⊇ *P*, is reconstructed. In this algorithm, *thresh* is a critical parameter for the accuracy of the tip reconstruction. Indeed, if noise is contained in the image profile, the reconstructed tip shape is strongly influenced by the geometries or dents of noise in the image profile. To choose the best *thresh* value, however, it is necessary to find a scale that well discriminates the heights of the sample surface geometry from the heights of the noise geometry, which is a difficult task, especially for novice users.

In the following, our differentiable BTR is introduced. The fact that the tip reconstruction is affected by noise in the original BTR can be regarded as an overfitting problem in machine learning. In machine learning, a typical approach to prevent overfitting is to use an appropriate loss functions and a regularization. The point of our idea is to statistically determine whether or not to carve the tip from the entire AFM data according to a loss function that takes noise into account, rather than individually determining whether or not to carve the tip from each dent in the image profile, as in the original BTR. Following the original BTR, our differentiable BTR is based on the relation *I* ∘ *P* = *I*, but the condition is relaxed because the equality does not hold in the presence of noise. Specifically, assuming that the noise is spatially independent white Gaussian noise, the loss function to be minimized would be,

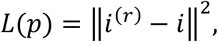

where ∥ ∥^2^ is the L2 norm. *i*^(*r*)^ is an image created by the opening *I*^(*r*)^ = *I* ∘ *P* = (*I* ⊖ *P*) ⊕ *P* with some tip shape *P*. Expanding the loss function by morphology operators, *i*^(*r*)^(*x, y*) can be rewritten as

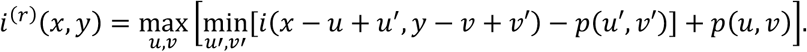

The differentiable BTR finds the optimal tip shape *p*(*u, v*) by minimizing this loss function over tip shape through the gradient of the loss function (Fig. 1e). Although max and min functions appeared in dilation and erosion seem to be non-differentiable at first glance, we treat this problem following the implementations of Max Pooling layer in convolutional neural networks (CNNs). For example, consider a one-dimensional tip shape *p*_*u*_ with *u* discretized by the index of a pixel. The gradient of *h* = max[*p*_1_, *p*_2_, *p*_3_] is, if *p*_3_ is the maximum, ∂*h*/∂*p*_1_ = 0, ∂*h*/∂*p*_2_ = 0, ∂*h*/∂*p*_3_ = 1, which is equivalent to argmax function. In image processing, some studies try to apply morphology operators to image processing by using argmax and argmin functions for differentiations^35^. In this study, we also use argmax and argmin functions for the differentiations of max and min functions, respectively.

Unfortunately, minimizing the above loss function is an ill-posed problem^36^. Other than the correct tip shape, for example, a very thin tip like the δ-function can make the value of the loss function almost zero (resulting in an overfit to noise for noisy AFM images). Therefore, regularization is introduced to find the upper bound on tip shape without an overfit while keeping the loss function small. In this study, we add the L2 norm of tip shape *P* as a regularization term,

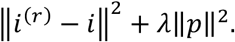

Here, *λ* is a weight parameter for the regularization term. When *λ* = 0, the minimization reduces to the original least-squares problem. When *λ* ≪ 1, the first term becomes dominant, and a sharp or acute tip shape might be chosen. As *λ* increases, the second term becomes dominant, and at some point, the overfit to noise is expected to be avoided and the reconstructed tip shape becomes blunt.

Moreover, we used a square pillar (*p*(*u, v*) = 0 for all (*u, v*) over the defined area for representing the tip) as an initial condition for the tip reconstruction, which is the same strategy as the original BTR. This choice of initial condition contributes to finding a blunt tip or the upper bound on tip shape. The optimization algorithm was implemented in a neural network/machine learning framework (the Flux.jl package^37^ in Julia programming language). Specifically, we implemented the dilation and the erosion as differentiable parts of neural network layers and optimized the tip shape in the same way in the training of neural networks (see Methods for details).

### Twin experiment: noise-free AFM images

By performing twin experiments, we compared the accuracies of two blind tip reconstruction algorithms, the original and end-to-end differentiable BTRs (Fig. 2). In the experiments, 20 frames of artificial pseudo-AFM images were generated from the structure of myosin V motor domain (PDB ID: 1OE9^38^) using a typical tip shape (a hemisphere combined with a circular frustum of a cone^20^, which serves as the ground truth tip). The reconstructed tip shape and molecular surface were compared to those of the ground truth (see Methods for details). For each pseudo-AFM image, the structure was randomly oriented. To understand the relationship between the parameters (*thresh* and λ) of the algorithms and the identity *I* ∘ *P* = *I*, we applied wide range of parameter values and monitored the loss function *L*(*p*) = ∥*i*^(*r*)^ − *i*∥^2^ and reconstructed tip shapes.

**Figure 2.**
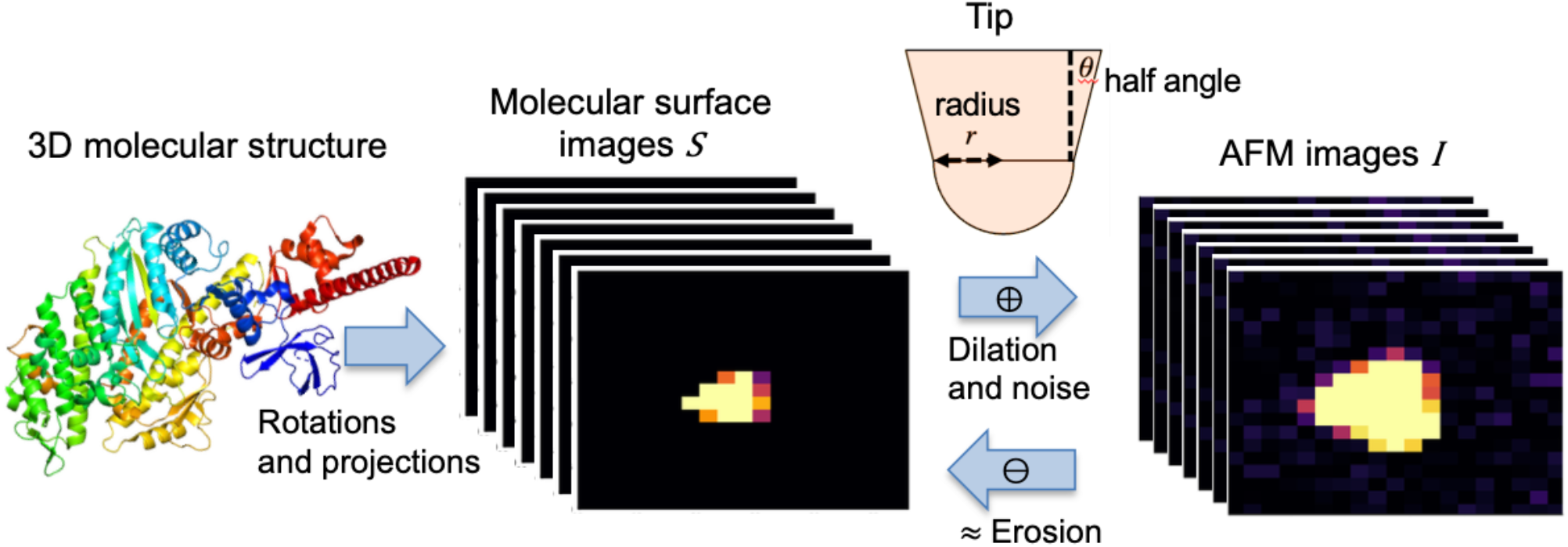
Schematic of twin experiments. From the structural model, the images of molecular surfaces are generated. Then, the molecular surfaces are converted to pseudo atomic force microscopy (AFM) images by dilation with a given tip shape. These molecular surfaces and tip shape are used as ground truths in the experiments.

Figure 3 shows the results of BTRs from noise-free AFM images. Under the noise-free condition, the original BTR can perfectly reconstruct the ground truth tip with *thresh* ≪ 1 (Fig. 3d left). *L*(*p*) increases as *thresh* increases (Fig. 3b), correlated with the thickness of the reconstructed tip shape (Fig. 3d right). The rate of the increase is slow for a range of small *thresh*, but is gradually accelerated later. The same is true for the differentiable BTR. The differentiable BTR also works perfectly to reconstruct the true tip shape with *λ* ≪ 1 (Fig. 3e left). *L*(*p*) increases as λ increases (Fig. 3c), reconstructing thicker tips (Fig. 3e right). Notably, the slope of the *L*(*p*) with respect to λ increases suddenly.

**Figure 3.**
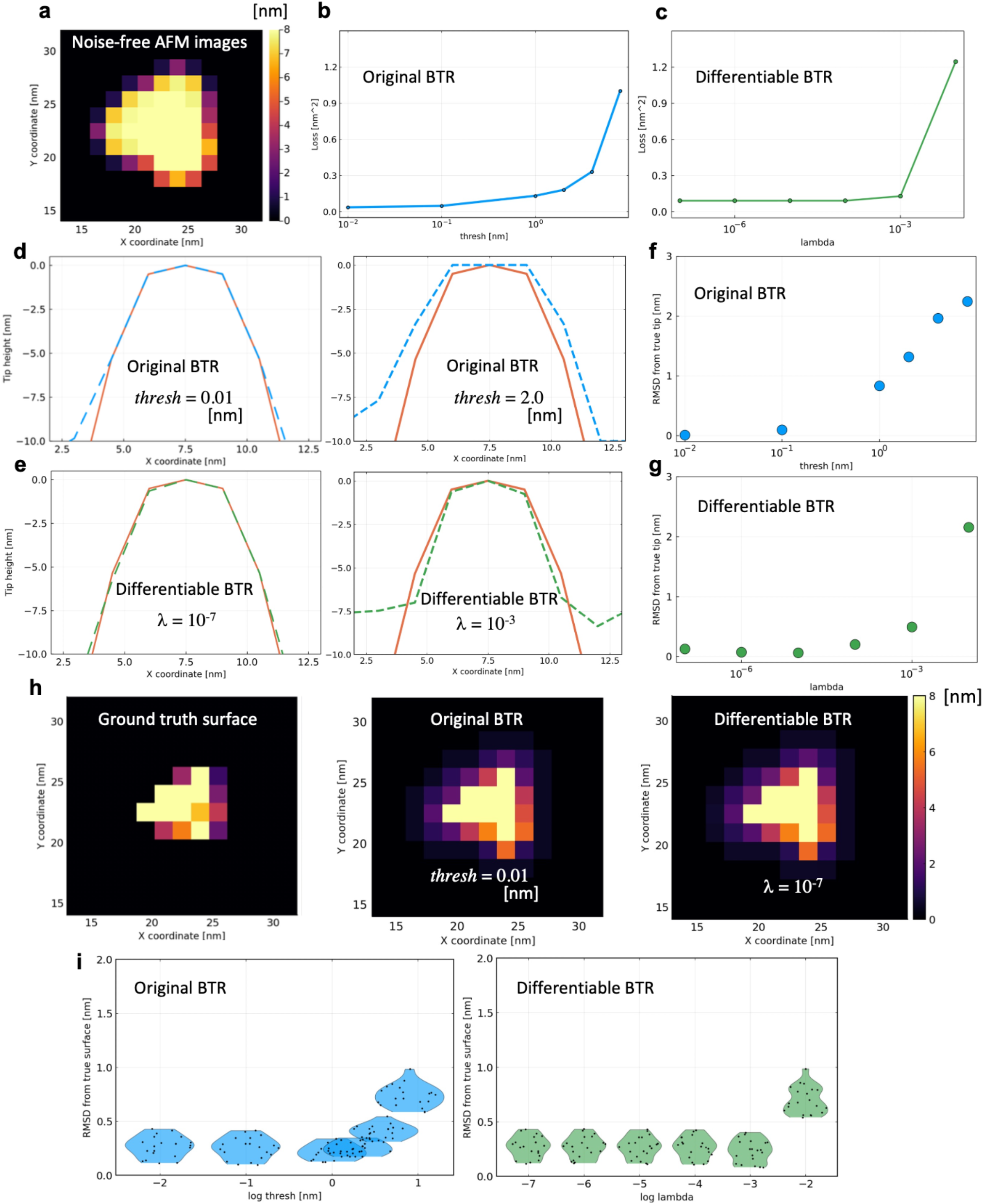
Results of twin experiment in noise-free condition. **a**, 1st frame of 20 images used for the twin experiment. **b-c**, Loss functions optimized at various parameter values. **d-e**, Cross sections of reconstructed tip shapes along the *x*-axis with the original and differentiable blind tip reconstruction (indicated by dashed blue lines and dashed green lines, respectively), compared with the ground truth (red line). **f-g**, Root mean square deviations (RMSDs) of the reconstructed tips from the ground truth (with the same coloring scheme). **h**, Reconstructed molecular surfaces by the deconvolutions with the reconstructed tips. **i**, RMSDs of the deconvoluted molecular surfaces of all 20 frames from the ground truth visualized by violin plots.

Figure 3h shows the molecular surfaces reconstructed by deconvolving the 1st frame of the pseudo-AFM images (Fig. 3a) of the data set with the reconstructed tips. Here, the deconvolution was performed by using the erosion. The results show the shape outlines of myosin V motor domain recovered by using both BTRs. Although the reconstructed molecular surfaces *S*^(*r*)^ by the erosion have a property of *S*^(*r*)^ ⊇ *S*, making the reconstructed surface thicker than the ground truth, the reconstructed molecular surfaces look qualitatively similar to that of myosin V motor domain for both BTRs. The accuracies of the reconstructed molecular surfaces are quantitatively evaluated by root mean square deviations (RMSDs) from the ground truth surfaces (Fig. 3i). In the calculation, deconvolutions were performed for all 20 frames of the images and compared with the ground truth. The RMSDs show that both BTRs have good accuracies for reconstructing molecular surfaces from noise-free AFM images.

### Twin experiment: noisy AFM images

We then compared the robustness of the two BTRs against noise by adding spatially independent Gaussian noise with a standard deviation of *σ* = 0.3 nm to the 20 frames of pseudo-AFM images used in the previous noise-free condition. Here, *σ* = 0.3 nm is a typical noise size in the current HS-AFM measurements^20^. 100 sets of noisy pseudo-AFM data (each containing 20 frames) were created using 100 different noise realizations. Again, to understand the relationship between the parameters (*thresh* and λ) and the identity *I* ∘ *P* = *I*, we applied parameter values in a wide range and monitored *L*(*p*) = ∥*i*^(*r*)^ − *i*∥^2^ (Figs. 4b and 4c). In both BTRs, *L*(*p*) sharply increases irrespective of noise realizations. However, when the tip shapes are plotted for the original BTR, reconstructed tip shapes seem to depend not only on parameters but also on noise realization (Fig. 4d). Indeed, even with the same *thresh* parameter 4.0 nm, the original BTR reconstructed the tip shape thinner or thicker than the ground truth, depending on the realization of the noise (Fig. 4d right). The tip shape tends to be thin for *thresh* ≪ 1 (Fig. 4d left). This means that the original BTR algorithm overfitted the tip shape to the noise in the pseudo-AFM images. The RMSD from the ground truth tip shows that *thresh* ≈ 4.0 nm is the best parameter value for accurate tip reconstruction (Fig. 4f). Parameter tuning of *thresh* has been discussed in previous studies. Villarrubia proposed, from the analysis of a single-frame pseudo-AFM image, to use 2-4 *σ* for thresh^1^. Although Villarrubia analyzed only a single-frame image, this value may depend on the number of frames used because the chance for noise level exceeding *thresh* changes by the number of samples. In this sense, *thresh* = 4.0 nm would be reasonable for the current 20-frame AFM images. Dongmo et al.^26^ proposed to monitor the change of tip volume by changing *thresh* parameter from low to high values. They discussed that the parameter where tip volume change becomes large would be regarded as the optimal one^26^. In the current context, that parameter value might correspond to the value where the loss function starts to increase sharply (*thresh* ≈ 4.0 nm in Fig. 4b).

**Figure 4.**
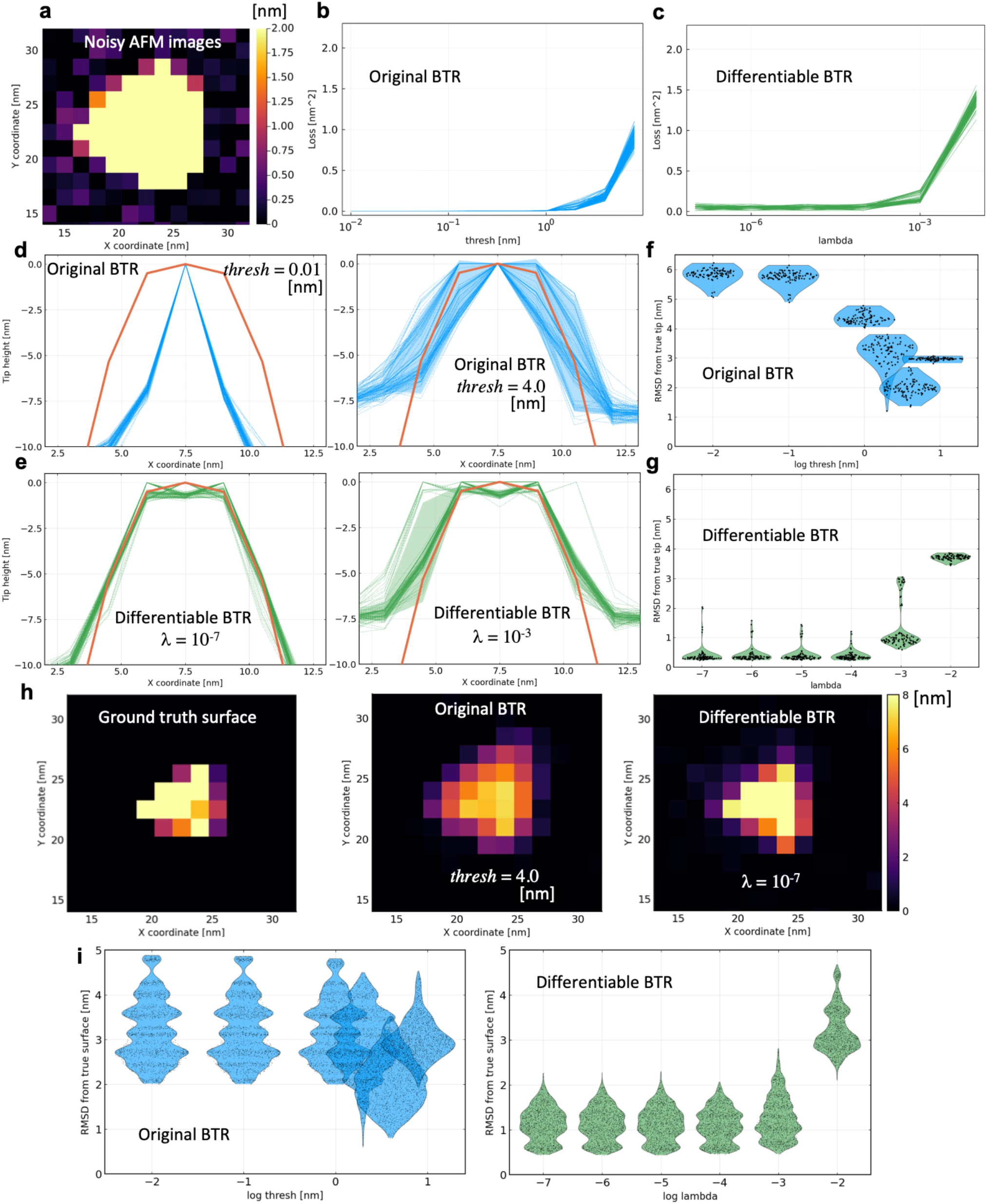
Results of twin experiment in noisy condition. **a**, 1st frame of 20 images used for the twin experiment. **b-c**, Loss functions optimized at various parameter values. **d-e**, Cross sections of reconstructed tip shapes along the *x*-axis with the original and differentiable blind tip reconstruction (indicated by dashed blue lines and dashed green lines, respectively), compared with the ground truth (red line). Shaded area represents the standard deviation. **f-g**, Root mean square deviations (RMSDs) of the reconstructed tips from the ground truth visualized by violin plots (with the same coloring scheme). **h**, Reconstructed molecular surfaces by the deconvolutions with the reconstructed tips. **i**, RMSDs of the deconvoluted molecular surfaces of all frames (20 images) from the ground truths visualized by violin plots.

On the other hand, the reconstructed tip shapes by the differentiable BTR show less dependence than the original BTR, and similar tip shapes (close to the ground truth) are reconstructed more stably for most noise realizations (Fig. 4e). Some tip shapes look distorted in the reconstructed tips of *λ* = 10^−3^ (Fig. 4e right), partially due to the boundary conditions of the tip image. Interestingly, while the tip shape became thicker as λ increases due to the regularization penalty (Fig. 4e right), the reconstructed tip shape did not fall into thin tip shapes even when the *λ* becomes small (*λ* ≪ 1, Fig. 4e left). This robustness in the differentiable BTR enables novice users to reproduce accurate reconstructions using a parameter value from a wide range. The average RMSD of the reconstructed tips from the ground truth is 2.0 nm for the original BTR with *thresh* = 4.0 nm, and 0.4 nm for the differentiable BTR with *λ* = 10^−7^ (Figs. 4f and 4g).

Figure 4h shows the molecular surfaces obtained by deconvolving the 1st frame of the pseudo-AFM images (Fig. 4a) using ones of the reconstructed tips. In Fig. 4h, the reconstructed molecular surface by the original BTR looks somewhat blurred because the erosion cannot “carve” the image profile sufficiently with a thin tip. On the other hand, the differentiable BTR successfully deconvolved the shape outlines of myosin V motor domain. The accuracies of the reconstructed surfaces were quantitatively evaluated by RMSDs from the ground truth (Fig. 4i). The surface reconstructions by the differentiable BTR outperform the original BTR in this noisy condition. The average RMSD of the reconstructed molecular surfaces from the ground truth is 2.1 nm for the original BTR with *thresh* = 4.0 nm, and 1.1 nm for the differentiable BTR with *λ* = 10^−4^.

### Twin experiment: double-tip effect

The double-tip effect is one of the most frequent artifacts in AFM measurements, which occurs when the tip is damaged or contaminated during the measurement^39^. Since AFM images measured by damaged or contaminated tips are often discarded without further analysis, it would be helpful if the double-shaped tip could be reconstructed only from AFM images and the doubled images were deconvolved to obtain true molecular surfaces. In this twin experiment, we mimicked the double tip effect with a double-peaked tip shape created by aligning the single tips used in the previous twin experiments along the *x*-axis. As in the previous twin experiment, we added spatially independent Gaussian noise with a standard deviation of *σ* = 0.3 nm to the pseudo-AFM images emulated by the double tips (Fig. 5a).

**Figure 5.**
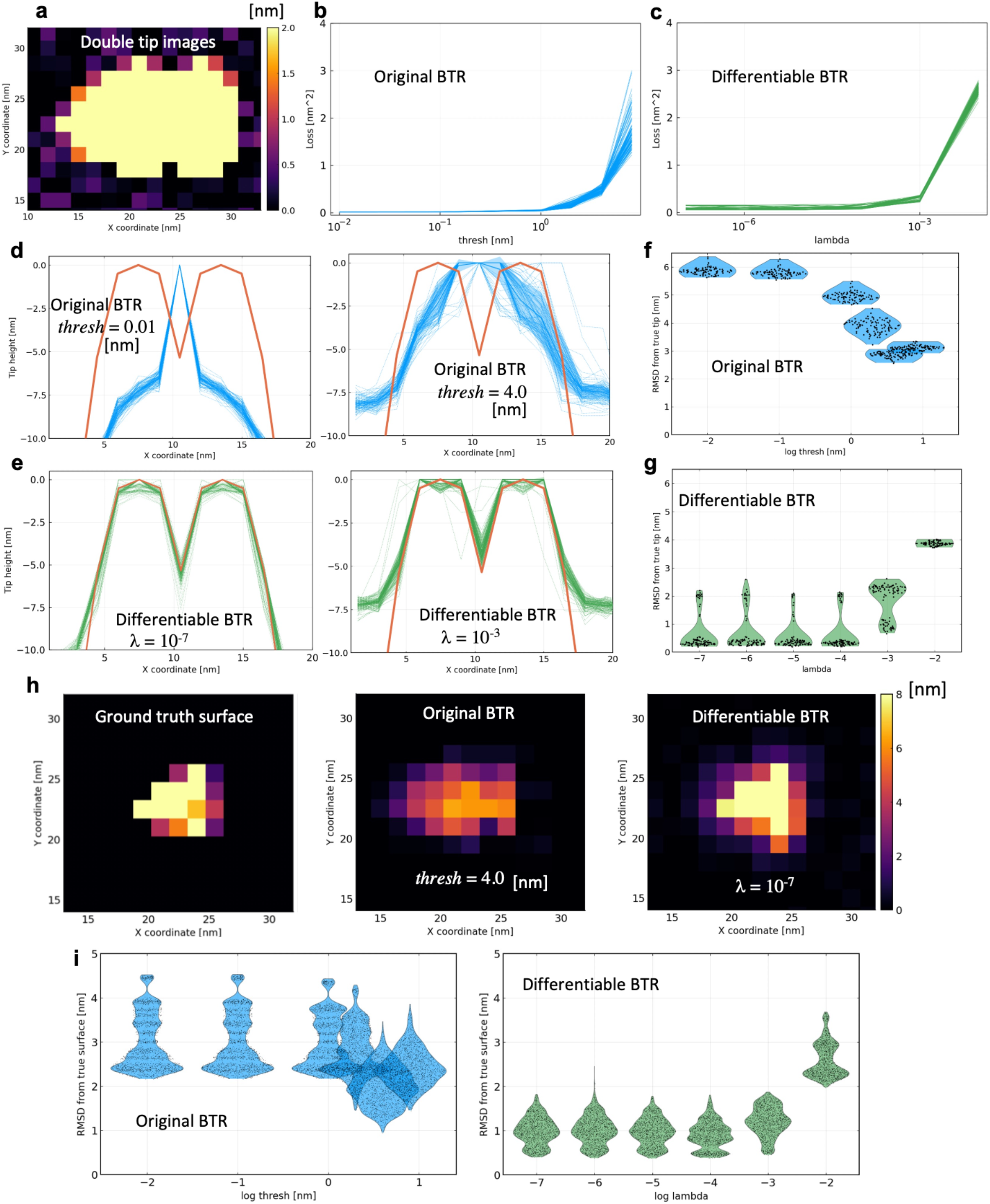
Results of twin experiment in a double-tip case. **a**, 1st frame of 20 images used for the twin experiment. **b-c**, Loss functions optimized at various parameter values. **d-e**, Cross sections of reconstructed tip shapes along the *x*-axis with the original and differentiable blind tip reconstruction (indicated by dashed blue lines and dashed green lines, respectively), compared with the ground truth (red line). **f-g**, Root mean square deviations (RMSDs) of the reconstructed tips from the ground truth visualized by violin plots (with the same coloring scheme). **h**, Reconstructed molecular surfaces by the deconvolutions with the reconstructed tips. **i**, RMSDs of the deconvoluted molecular surfaces of all frames (20 images) from the ground truths visualized by violin plots.

Figure 5d shows tip shapes reconstructed with the original BTR. The original BTR failed to detect double tips, and the reconstruction resulted in single-peak tips for most parameter values. On the other hand, the differentiable BTR successfully detected double tips, and their shapes are roughly consistent with the ground truth over a wide parameter range (Fig. 5e). Again, translational shifts in some of the reconstructed tips are observed due to the boundary effect of the tip image. The result does not mean that the original BTR algorithm inherently cannot detect double tips. Indeed, under noise-free condition, the original BTR is able to reconstruct the double-tip shape with high accuracy. In the original BTR algorithm, the tip apex is set to be at the origin of the *uv*-coordinates, and the apex should be fixed at the origin during the reconstruction without any modifications. In noisy conditions, while one of two tip apexes is fixed at the origin, the other tip apex is irreversibly “carved” by chance due to noise, making it difficult to detect double tips. On the other hand, since the algorithm of the differentiable BTR can reconstruct the tip without imposing any constraints on the tip apex, it can stably detect double tips. In the differentiable BTR, the origin of the *uv*-coordinates is set to the center of mass of the tip (see Methods for details). The average RMSD of the reconstructed tips from the ground truth is 2.9 nm for the original BTR with *thresh* = 4.0 nm, and 0.5 nm for the differentiable BTR with *λ* = 10^−5^ (Figs. 5f and 5g).

Figure 5h shows the molecular surfaces obtained by deconvolving the 1st frame of the pseudo-AFM images (Fig. 5a) using one of the reconstructed tips (Figs. 5d and 5e). With a single tip reconstructed by the original BTR, the double-tip artifact remains in the molecular surface even after the deconvolution, and the molecular shape looks doubled. On the other hand, double tips reconstructed with the differentiable BTR can remove the doubled molecular shapes and reconstruct a molecular surface much closer to the ground truth. The accuracies of the reconstructed surfaces were quantitatively evaluated by RMSDs from the ground truth ones (Fig. 5i). The average RMSD of the reconstructed molecular surfaces from the ground truth is 2.0 nm for the original BTR with *thresh* = 4.0 nm, and 0.9 nm for the differentiable BTR with *λ* = 10^−4^.

### Real experimental data: Myosin V walking along actin filament

Finally, we compared the two BTR algorithms by analyzing real AFM data. We analyzed the HS-AFM data of myosin V measured by Kodera et al^5^. Myosin V is a homo-dimeric molecular motor that moves linearly along the actin filament driven by ATP hydrolysis free energy. Myosin V serves to transport cargos attached to the center of the dimer. The HS-AFM movies by Kodera et al successfully capture the events that myosin V walks on the actin filament coupled with its ATP-dependent conformational change. It is important to deconvolve the AFM images to obtain the detailed shape of the Myosin V’s molecular surface during the walking step. Here, we analyzed 30 frames of a HS-AFM movie that capture a single walking step of Myosin V.

As illustrated in Fig. 6a, the HS-AFM images contain numerous horizontal scars emanating from the molecules to the right direction along the *x*-axis, which corresponds to the scanning direction of the tip. These scars are often called the parachuting artifact^40^. The “parachuting” means that, when the tip scan velocity is high and the target molecular height suddenly decreases along the tip scanning line, the tip can completely detach from the molecular surface. It takes some time for the tip to land on the stage or the sample surface at a different point, creating scars in the direction of scanning^40^. The parachuting artifact appears when the tip scan speed is fast, as in some HS-AFM measurements. Since scars caused by the parachuting artifact are not stochastic but deterministic, we recognize the scars as part of the signal and try to reflect them in the effective tip shape.

**Figure 6.**
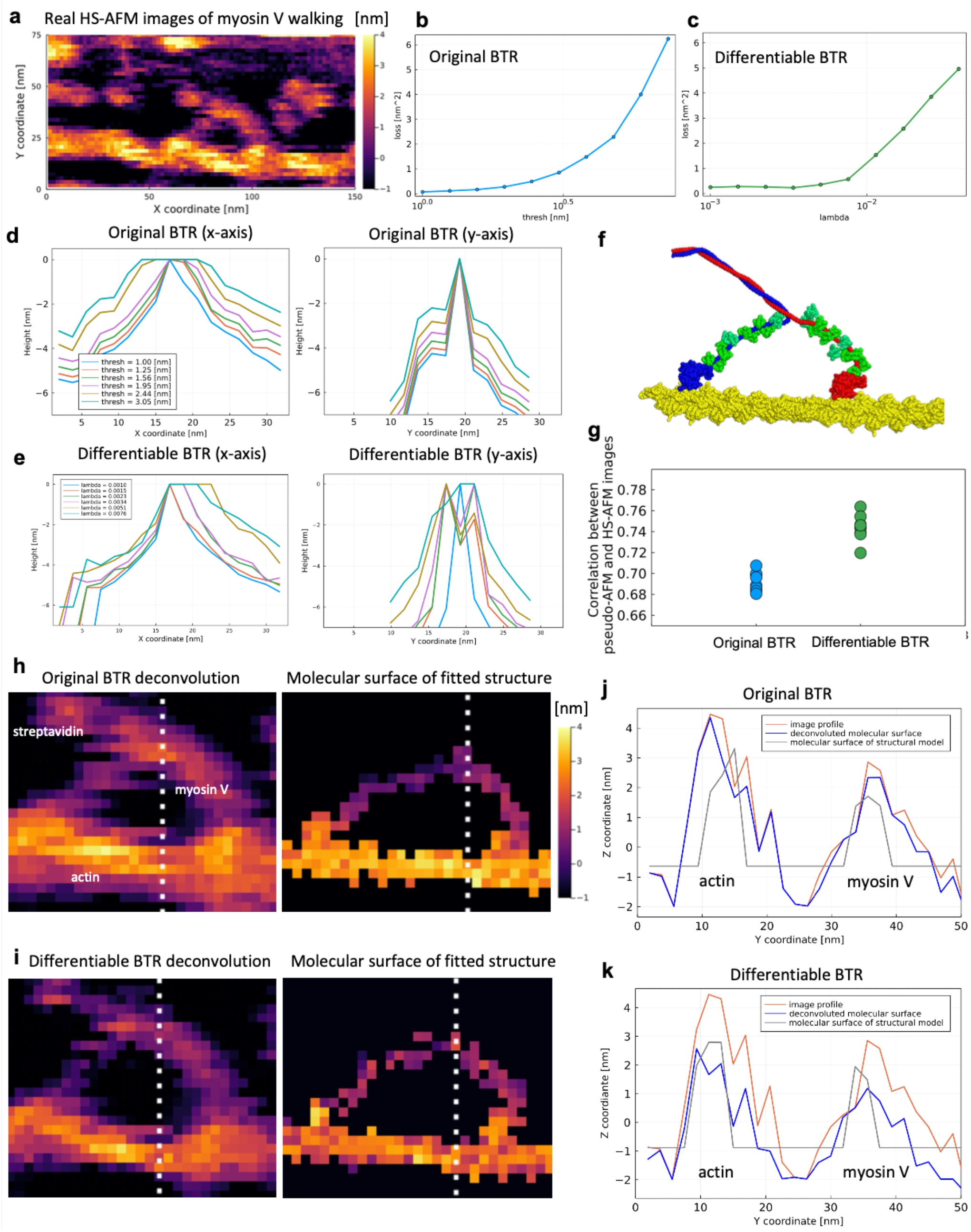
Blind tip reconstruction (BTR) from the high-speed atomic force microscopy (HS-AFM) data of myosin V walking. **a**, 27th frame of 30 images used for the analysis. **b-c**, Loss functions optimized at various parameter values. **d-e**, Cross sections of reconstructed tip shapes along the *x*-axis with the original and differentiable BTR. **f**, Representative structure of 10 structures taken from previous flexible fitting molecular dynamics simulations. **g**, Maximum correlation coefficients between the 27th HS-AFM image and pseudo-AFM images generated from each of 10 structures. **h**, Reconstructed molecular surfaces created by deconvolutions with the BTR tips compared with the molecular surfaces of structured models with the maximum correlation coefficients. The while dashed lines indicate the cross sections used in (**j**) and (**k**). **j-k**, Cross sections of the molecular surfaces along the *y*-axis. Red lines are the image profiles, blue lines are the reconstructed molecular surfaces, and gray lines are the molecular surfaces created from the structural models.

Figures 6d and 6e show the *x*- and *y*-axes cross sections of the tip shapes reconstructed from the real HS-AFM images by the original and differentiable BTRs for various *thresh* and λ parameters. The tip shapes reconstructed with the original BTR tend to be thicker along the *x*-axis (Fig. 6d left) while very thin along the *y*-axis (Fig. 6d right). This is largely due to the scars of the parachuting artifact. The images along the parachuting lines, i.e., along the *x*-axis, are deterministic signals. However, along the *y*-axis, whether the parachuting appears or not are quite stochastic: The parachuting appears only when the fluctuating molecular surface happens to be high (large *z*-coordinates). Therefore, along the *y*-axis, dents caused by the differences between parachuting and non-parachuting lines should be recognized as stochastic noise. The original BTR misinterpreted that the large dents along the *y*-axis can be ascribed to the tip shape and excessively “carved” the tip shape. While the tip shapes reconstructed by the differentiable BTR well capture a smooth attenuation by parachuting on the right side of the *x*-axis (Fig. 6e left), those along the *y*-axis do not become thin as in the original BTR but maintain a certain degree of thickness with less parameter dependence (except for a very small weight, *λ* = 0.0010, Fig. 6e right).

Figures 6h and 6i show the molecular surfaces of the 27th frame of the HS-AFM movie deconvolved with the reconstructed tip shapes by the original and differentiable BTRs, respectively. The 27th frame was chosen because flexible fitting molecular dynamics simulations were performed for this frame in our previous study^41^. For the differentiable BTR, we used λ = 0.0051 for the deconvolution, which corresponds to the value where the loss function starts to increase sharply (following the parameter tuning scheme by Dongmo et al.^26^) (Fig. 6c). For the original BTR, we used the tip of *thresh* = 1.95 nm because the tip apex radius of this parameter value is comparable to (or slightly larger than) that of the differentiable BTR with the chosen λ = 0.0051 (see Figs. 6d and 6e). In the molecular surface deconvolved by the tip of the original BTR (Fig. 6h), the legs of Myosin V are thinner than in the image profile, but there are still horizontal scars due to the parachuting artifact. On the other hand, the molecular surface deconvolved by the tip shape of the differentiable BTR has fewer parachuting lines (Fig. 6i). This is because the differentiable BTR has a thicker tip shape along the *y*-axis direction, which contributes to more extensive “carvings” by the erosion of the image profiles along the *y*-axis. The same trend can be seen in all the frames (Supplemental Movie1).

Are the molecular surfaces obtained by the deconvolutions accurate? To verify this, we compared these surfaces with structural models. Here we used 10 structures taken from our previous flexible fitting molecular dynamics simulation study of the 27th frame of the same data^41^. In the study, the structures of myosin V and actin filament were modeled using the crystal structure by homology modeling, and then coarse-grained molecular dynamics simulations were performed, imposing restraints on the structure fitted to the 27th frame of the experimental HS-AFM movie. Since the tip shape used in the flexible fitting is not the same as the current study, we selected 10 plausible structures as templates to make a comparison with the deconvoluted surfaces (Fig. 6f). Each structure was exhaustively translated and rotated to create pseudo-AFM images, and the correlation coefficients of those images with the target frame of the HS-AFM data were calculated. We recorded the maximum correlation coefficients and corresponding poses (translation and rotation) for the 10 structures. Figure 6g compares the maximum correlation coefficients of the 10 structures for both BTRs. The figures suggest that the differentiable BTR reproduces pseudo-AFM images more similar to the real HS-AFM image. This further means that the tip shape is consistent with structural models and the HS-AFM data. Figures 6j and 6k show the cross sections of the deconvoluted molecular surfaces and the surfaces obtained by the structural models that have the maximum correlation coefficients for two BTRs, respectively. In the figure, actin filament is located at around *y* = 13 nm. We can see that the deconvolved surface with the original BTR is still thicker than the surface of the structural model. In contrast, the surface of the differentiable BTR is consistent with the structural model. Again, this result implies the accuracy of the tip reconstructed by the differentiable BTR.

## Discussion

In this study, we have proposed the end-to-end differentiable BTR method based on a loss function and a regularization term considering noise in AFM image to avoid overfitting. The loss function is systematically optimized following the framework developed by the recent advances in deep learning studies. The results of the twin experiments showed that the differentiable BTR is more robust to noise than the original BTR. Finally, demonstration of real HS-AFM data shows that the differentiable BTR reconstructs tip shapes and the deconvoluted molecular surface is consistent with the structural model. Since the method is a quite general, we expect that the method would become a routine method for analyzing noisy AFM images as well as HS-AFM data.

The twin experiments have shown that reconstructed tip shapes with the differentiable BTR are less dependent on the parameter *λ*. For relatively small noise (*σ* < 0.3 nm), any small *λ* values (*λ* ≪ 1) are expected to reproduce tip shapes close to the ground truth one. For relatively large noise (*σ* > 1.0 nm), as seen in the real HS-AFM data of myosin V walking, small *λ* may reconstruct thin tip shapes even with the differentiable BTR. In such cases, choosing *λ* values where the loss function starts to increase rapidly is recommended to reconstruct reasonably thick tip shapes. As with the case of neural network frameworks, there are several hyperparameters in the optimization process in the differentiable BTR. In particular, the number of epochs should be carefully chosen. If the epoch number is too large, that increases the chance to overfit to noise, while too small epochs result in insufficient learning. Using techniques such as early stopping, or hyperparameter optimization tools^42^, may work to determine an appropriate epoch number.

As mentioned in Introduction, Tian et al. proposed an improved regularization scheme against noise in the framework of original BTR^27^, and Jóźwiak et al. simplified Tian’s idea to the case of standard AFM tips^28^. In this scheme, the update procedure of tip shape becomes more conservative, and it is thus expected to be robust against noise, as shown by Jóźwiak et al.^28^ Supplementary Fig. 1 shows the results of the original BTR using this regularization scheme for noisy single-tip and double-tip twin experiments. As expected, the regularization scheme improves the estimation and reconstructs blunter tip shapes compared to the original BTR. However, the RMSDs from the ground truth tip shape and molecular surface show that the differentiable BTR still outperforms the original BTR with the regularization scheme.

Since this study mainly targets the analysis of HS-AFM data, pseudo and real AFM data consisting of 20-30 frames were mainly analyzed. To check the dependence of reconstruction accuracy on the number of frames, we conducted noisy single-tip twin experiments using 1, 10, and 100 pseudo-AFM frames (Supplementary Fig. 2). Although the accuracies of tip and surface reconstructions are comparable in both BTRs for a single frame, the differentiable BTR outperforms the original BTR for 10 and 100 frames. An only drawback of the differentiable BTR here is that the computation time is much longer than the original BTR, especially for the analysis of 100 frames. Implementing GPU kernels for dilation and erosion, or using smooth functions instead of max and min, would accelerate the convergence of the optimization.

A concern with differentiable BTR is the relationship between ground truth tip shape and erosion. To reconstruct the tip accurately with differentiable BTR, erosion must be a good approximation to the inverse function of dilation. If the ground truth tip has a cone shape, the half angle must be small for the erosion to approximate the inverse function of the dilation well. As the half angle increases and exceeds a threshold value (e.g., 30 degrees), the accuracy of tip reconstruction deteriorates quickly because erosion fails to approximate the inverse function. However, as shown by the numerical study by Sumikama et al.^43^, in the tapping mode of HS-AFM, the half angle is expected to become rarely large because the effective tip shape becomes thinner. Also, one possible direction to overcome this problem would be to employ a CNN as a substitute for erosion. If the amount of data is large enough, a CNN is expected to approximate the inverse function of dilation well. In the analysis of the HS-AFM data of Myosin V, we reconstructed the effective tip shape regarding the scars caused by the parachuting effect as a signal. However, there may be cases where we want to estimate the real physical tip shape after removing the parachuting effect. In this case, it may be possible to divide the dilation into two layers in the differentiable BTR. The first dilation introduces a tip shape that adds lines linearly on the right side of the molecule, and the slope of the lines is parametrized. The second dilation is a standard dilation with a tip shape *p*(*u, v*). In such a framework, it would be possible to estimate both parameters simultaneously from AFM images. Since dilation is implemented as a part of the neural network layers, it is straightforward to extend the current BTR algorithm in this way. From another perspective, the loss function can be changed according to the types of AFM data. For examples, if AFM data contain large noise as outliers, using the L1 norm instead of the mean squared error (used in this work) for the loss function may work better for tip reconstruction.

Recently, Heath et al. developed localization AFM (LAFM)^44^, an elegant image reconstruction technique to overcome resolution limitations in AFM. In this approach, peak positions contacted with the tip apex in AFM image profiles are detected and accumulated to increase the resolution. Although the LAFM currently only uses the peak positions in the localization algorithm, it might be possible to accumulate positions contacted with the side of the tip for the localization algorithm if the tip shape is accurately reconstructed with the differentiable BTR in the future.

## Methods

### Original blind tip reconstruction

The C language source codes provided by Villarrubia^1^ were ported to the Julia programming language. Inside the codes, *xy*-coordinates and *uv*-coordinates are discretized by pixel indices (*x, y*) and (*u, v*), respectively. Starting with a square pillar tip, i.e., *p*_*u,v*_ = 0 (for any (*u, v*), zero is the maximum height for the inverted tip) as an initial condition, the tip shape was iteratively “carved” according to the dents in neighboring pixels until no “carving” events occur. For the analysis of multiple frames, the algorithm was sequentially applied to frames as proposed by Villarrubia^1^.

### Differentiable blind tip reconstruction

We implemented the differentiable BTR in Julia programming language. The *xy*-coordinate and *uv*-coordinate coordinates were discretized and used as pixel indices. The following algorithm was designed to minimize the loss function 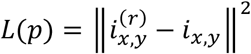 with respect to *p*_*u,v*_:

1. Initialize the tip shape to be a square pillar shape, *p*_*u,v*_ = 0 (for any *u* and *v*). Here, zero is the maximum height for the inverted tip.
2. Calculate the gradient of the loss function 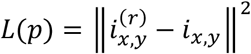
3. Update *p*_*u,v*_ according to the computed gradient using AdamW^45^ as an optimizer imposing a regularization term of the L2 norm ∥*p*_*u,v*_∥^2^. Learning rate of 0.1 nm, exponential decay for the first *β*_1_ of 0.9 and the second *β*, of 0.999 were used.
4. Trim values greater than zero. Positive *p*_*u,v*_’s are changed to zero, *p*_*u,v*_ = 0.
5. Using the height of *p*_*u,v*_ relative to the minimum height of *p*_*u,v*_ as weights, translate *p*_*u,v*_ so that its center of weight is located at the origin of the *uv*-coordinate. For the unknown *p*_*u,v*_ near the boundary (edge) created due to the translation, the minimum value of *p*_*u,v*_ was assigned.
6. Repeat step 2 to step 5 using a batch size of 1 image until 100-2000 epochs.

Dilation and erosion were implemented as layers of the neural network framework (the Flux.jl package^37^ in Julia language), and their custom pullbacks were defined using the ChainRulesCore.jl package^46^. Differentiation of opening and the backpropagation of the loss function were computed with the automatic differentiation by the package Zygote.jl^47^. The gradient of each dilation and erosion was tested during the development.

### Twin experiments

To evaluate the accuracy of the tip shape reconstruction, we performed twin experiments. We first generated pseudo-AFM images from a structure of myosin V motor domain without bound nucleotide (PDB ID: 1OE9^38^) using a specific tip shape, which serves as a ground-truth tip shape. The light chains were removed from the original crystal structure. Using the pseudo-AFM images, we performed the BTR and compared the tip shape reconstructed from the images with the ground truth tip shape. We set the *z*-axis of the *xy*-plane system to be the direction orthogonal to the stage (i.e., the *xy*-plane is parallel to the stage). Generated pseudo-AFM images consist of 30 × 30 pixels, where the width and height of each pixel are both 1.5 nm. In the simulation, the center of mass of the motor domain was placed at the origin of the *xy*-coordinate. Then the motor domain was randomly rotated in three dimensions. The heights of the van der Waals (vdW) sphere surfaces at the *xy*-coordinates at the pixel centers were calculated for neighboring atoms of each center, and the largest height was taken as the molecular surface. After obtaining the molecular surface at the pixel centers, the tip shape was used to create a pseudo-AFM image by performing dilation. We used the pixel size of 10 × 10 for the single-tip, 14 × 10 for the double-tip, and 17 × 11 for the HS-AFM data of myosin V walking. Finally, spatially independent Gaussian noise with a standard deviation of 0.3 nm was added to each pixel to obtain noisy pseudo-AFM images. The standard deviation of 0.3 nm is a typical noise size for the current HS-AFM^20^. For the single tip twin experiment, the tip shape of a hemisphere (with radius *r*) combined with a circular frustum of a cone, its apex radius of 2.5 nm and half angle of 10 degrees were used as the ground truth. For the double-tip twin experiment, two single tips were aligned in the *x*-axis direction. By repeating the above procedure, 20 pseudo-AFM images were simulated for each setup.

In the calculation of the root mean square deviation (RMSD) from the ground truth tip shape, *uv*--coordinates where the ground tip *p*(*u, v*) > −7.0 nm were used. 7.0 nm roughly corresponds to the height of myosin V motor domain. In the calculation of the RMSD from the ground truth molecular surface, *xy*-coordinates where the surface height *s*^(*r*)^(*x, y*) > 1.0 were used. 1.0 nm roughly corresponds to 3*σ* of noise. Considering translational shifts in the reconstructed tips, we translated the tips exhaustively and recorded the minimum RMSD value for each tip.

### HS-AFM data of myosin V

HS-AFM measurements of myosin V walking along actin filaments were carried out by Kodera et al., and details for the experiments are given in the paper^5^. In the measurements, the ATP-driven translocations of Myosin V (M5-HMM) along actin filaments were observed by the HS-AFM. Actin filaments were partially biotinylated and immobilized using streptavidin on biotin-containing lipid bilayers formed on a mica surface (stage). A positively charged lipid was included in the bilayer to facilitate the weak sideways adsorption of M5-HMM onto the bilayer surface. HS-AFM measurements were conducted with the tapping mode. This study used 30 frames of HS-AFM data sets, which well captured single-step walks of M5-HMM. Each image consists of 80 x 40 pixels, and the resolution of each pixel is 1.875 nm x 1.875 nm. We used these 30 frames to perform original and differentiable BTRs over various parameters. The tilt of the stage in the HS-AFM data was corrected by fitting a *xy*-plane to the data.

### Analysis of HS-AFM data

Structural modeling and flexible fitting molecular dynamics simulations of Myosin V and actin filament were performed in the same way as in the previous study by Fuchigami and Takada^41^. First, we generated all-atom models of a tail-truncated myosin V (each of two heads is in the lever-arm down and up conformations) and an actin 31-mer filament using MODELLER version 9.25^48^. Then, we constructed coarse-grained models of the actin-myosin complex in the down-down state using CafeMol version 3.2.0^49^. We performed ten flexible fitting simulations of the actin-myosin complex in the down-down state using structure-based AICG2+ potential^50^, AFM stage potential, and AFM image-based bias potential^51^. See the reference^41^ for more details.

Using ten structures with the highest cosine similarity in each of the ten flexible fitting simulations, we performed the rigid-body fitting of 3D structures to 2D HS-AFM images of Myosin V. For each structure model, we exhaustively rotated the structure around the rotation axis (roll axis) of the helix of actin filament by 3 degrees (within the range of 60 degrees), combined with the rotation around the yaw axis of the filament by 3 degrees (within the range from −30 to 30 degrees starting from the actin filament parallel to the *x*-axis). For all the combinations of two angles, we created pseudo-AFM images from the structure. The pseudo-AFM images were created in the same way as in the twin experiment. For the radii of coarse-grained Cα beads, we used the effective radius of amino acid residues used in CryoEM data analysis^52^. The coiled coil domain of myosin V were ignored in the generation of the pseudo-AFM images because this domain was no clearly observed in the HS-AFM. Since the height scale in the real HS-AFM data roughly corresponds to half of the structural model, we doubled the height of the tips reconstructed from the real HS-AFM images. Using it, we then applied the dilation to the structure-model molecular surface. The pseudo-AFM images from the structure model were translated over possible locations on the real HS-AFM image (note that the pseudo-AFM image is smaller than the HS-AFM images because it contains only a single myosin V and a fragment of actin filament), and we computed the correlation coefficient between the overlapped regions of two images. We recorded the maximum correlation coefficient between the pseudo-AFM image and the real AFM image and its corresponding pose (translation and rotation). The structure, of which pseudo-AFM image has the maximum correlation coefficient, was compared with the molecular surface obtained by the deconvolution of the HS-AFM image. In the comparison, the height of the molecular surface of the structural model was transformed by 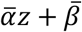, where 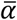 and 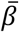 were determined by the least squares fitting to the deconvoluted surface. This transformation was needed to adjust the calibration and the offset level.

## Data availability

Jupyter notebooks to reproduce the results of this paper are publicly available at https://github.com/matsunagalab/differentiable_BTR. Some notebooks automatically download the structure of the target molecule from Protein Data Bank and calculate pseudo-AFM images used in twin experiments. The structural models of myosin V and actin filament are included in the repository. Real HS-AFM data of myosin V walking along actin filament are available upon reasonable request and with the permission of Kodera et al^5^.

## Code availability

Jupyter notebooks for this study to reproduce the results are publicly available at a GitHub repository, https://github.com/matsunagalab/differentiable_BTR. The basic functions including dilation, erosion, and their custom pullbacks are implemented in our in-house Julia package, MDToolbox.jl, which is publicly available at https://github.com/matsunagalab/MDToolbox.jl.

## Acknowledgement

We thank Noriyuki Kodera for giving the HS-AFM data of myosin V walking and suggesting us to apply the method to a double tip case. We thank Tohru Niina for stimulating discussion on the blind tip reconstruction. We also thank Ryuhei Oshima for his help to prepare Fig. 1. This work was supported by JST CREST (Grant number: JPMJCR1762 to Y.M. and S.T.), MEXT as “Program for Promoting Researches on the Supercomputer Fugaku” (Biomolecular dynamics in a living cell, Grant number: JPMXP1020200101 to Y.M. and S.T.), JSPS KAKENHI (Grant number: 20K21380 to Y.M.), and the Cooperative Research Program of “Network Joint Research Center for Materials and Devices” (to Y.M.).

---

**Supplementary Figure 1.**
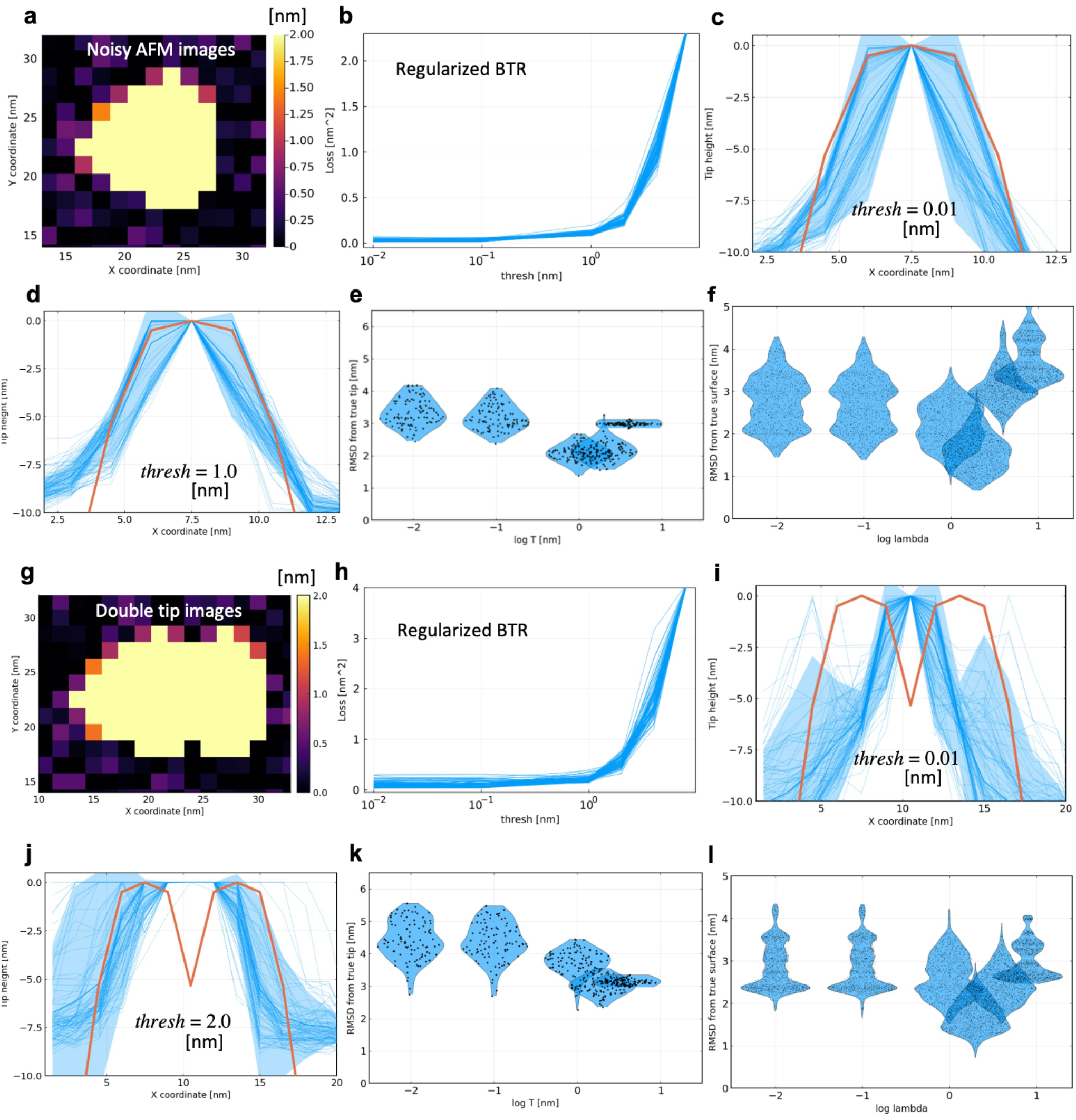
Results of twin experiments for the original blind tip reconstruction with a improved regularization scheme. **a**, 1st frame of 20 images used for the twin experiment in noisy conditions. **b**, Loss functions optimized at various parameter values. **c-d**, Cross sections of reconstructed tip shapes along the *x*-axis (indicated by dashed blue lines), compared with the ground truth (red line). Shaded area represents the standard deviation. **e**, Root mean square deviations (RMSDs) of the reconstructed tips from the ground truth visualized by violin plots. **f**, Reconstructed molecular surfaces by the deconvolutions with the reconstructed tips. **g-l**, Results of twin experiment in a double-tip case.

**Supplementary Figure 2.**
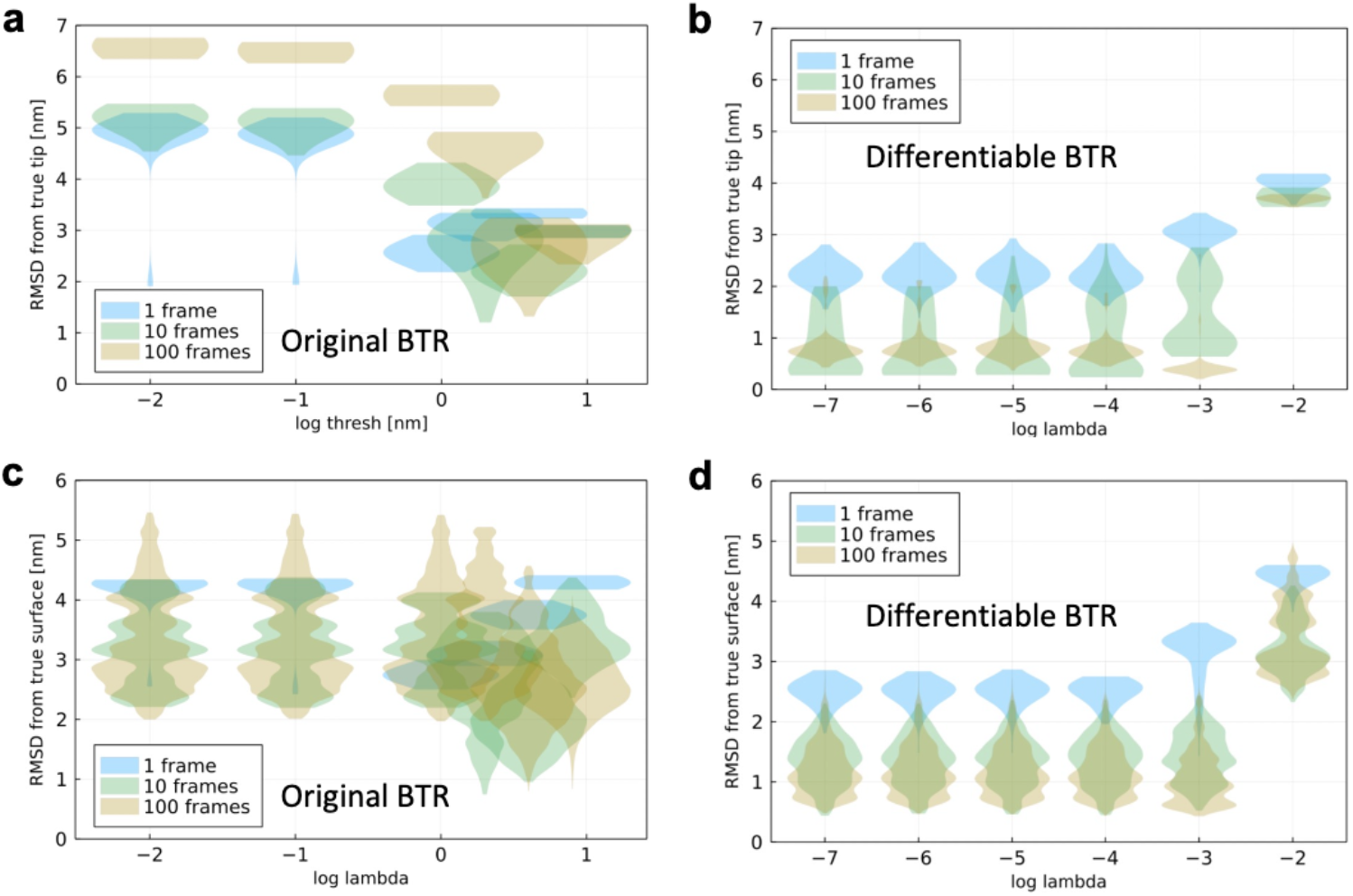
Results of twin experiment in single-tip noisy conditions with various number of frames (1 frame, 10 frames, and 100 frames). **a**, Root mean square deviations (RMSDs) of the reconstructed tips with the original blind tip reconstruction (BTR) from the ground truth visualized by violin plots. **b**, RMSDs of the reconstructed tips with the differentiable BTR from the ground truth. **c**, RMSDs of the reconstructed molecular surface with the original BTR from the ground truth. **d**, RMSDs of the reconstructed molecular surface with the differentiable BTR from the ground truth.

**Supplementary Movie 1**. Reconstructed molecular surfaces of myosin V and actin filament by the deconvolutions with the tips of blind tip reconstructions. The whole frames (30 frames) are converted to a movie of 5 fps.

## Notes

### Competing Interest Statement

The authors have declared no competing interest.

